# Temporal Stability and Molecular Persistence of the Bone Marrow Plasma Cell Antibody Repertoire

**DOI:** 10.1101/066878

**Authors:** Gabriel C. Wu, Nai-Kong V. Cheung, George Georgiou, Edward M. Marcotte, Gregory C. Ippolito

**Affiliations:** Center for Systems and Synthetic Biology University of Texas at Austin, Austin, TX, USA; Department of Molecular Biosciences, University of Texas at Austin, Austin, Texas, USA; Department of Pediatrics, Memorial Sloan-Kettering Cancer Center, New York, New York, USA; Department of Biomedical Engineering, University of Texas at Austin, Austin, Texas, USA; Department of Chemical Engineering, University of Texas at Austin, Austin, Texas, USA; Institute for Cell and Molecular Biology, University of Texas at Austin, Austin, TX, USA

## Abstract

Plasma cells in human bone marrow (BM PCs) are thought to be intrinsically long-lived and to be responsible for sustaining lifelong immunity through the constitutive secretion of antibody—but the underlying basis for this serological memory remains controversial. Here, we analyzed the molecular persistence of serological immunity by an examination of BM PC immunoglobulin heavy-chain (IGH) transcripts derived from serial bone marrow specimens obtained during a span of several years. Using high-throughput sequence analysis of the same individual for 6.5 years, we show that the BM PC repertoire is remarkably stable over time. We find that the bias in IGH V, D, and J individual gene usage and also the combinatorial V–D, V–J, D–J, and V-D-J usage across time to be nearly static. When compared to a second donor with time points 2 years apart, these overall patterns are preserved, and surprisingly, we find high correlation of gene usage between the two donors. Lastly, we report the persistence of numerous BM PC clonal clusters (~2%) identifiable across 6.5 years at all time points assayed, supporting a model of serological memory based, at least in part, upon intrinsic longevity of human PCs. We anticipate that this longitudinal study will facilitate the ability to differentiate between healthy and diseased antibody repertoire states, by serving as a point of comparison with future deep-sequencing studies involving immune intervention.

## INTRODUCTION

The human bone marrow (BM) is a specialized immune compartment that is responsible for both the initial generation of newly-formed B cells and also the maintenance of terminally differentiated, antibody-secreting plasma cells (PCs). The BM, and the PCs it harbors, is a major site of antibody production and is the major source of all classes and subclasses of human immunoglobulins (Ig) detectable in the serum^1,2^. lg-secreting BM PCs are generally believed to be “long-lived” and maintained for the lifespan of the organism^3^. In this regard, it has been well established in longitudinal serological studies that antiviral serum antibodies can be remarkably stable, with halflives ranging from 50 years (e.g. varicella-zoster virus) to 200 years for other viruses (e.g. measles and mumps); however, in contrast, antibody responses to non-replicating antigens (e.g. tetanus and diphtheria bacterial toxins) rapidly decay with much shorter half-lives of only 10–20 years^4^. Not only does this suggest that antigen-specific mechanisms play a substantial role in the establishment and/or maintenance of serological memory, but raises the question of whether the differential stability of antibody responses might reflect differential intrinsic longevity of PCs themselves. This has been previously proposed in the context of vaccinations and infections^4,5^, and is also supported by observations of differential stability of autoantibody titers when using B-cell depleting therapies to treat autoimmune diseases^6,7^.

The basis underlying lifelong serological memory (antibody responses) remains controversial^3,8,9^. Data supporting a model for intrinsic longevity in PC survival (and hence longevity in serum antibody maintenance) has been posited for the laboratory mouse^10,11^, but data for human PCs are absent. Based upon the murine models, it has been assumed that human BM PCs are similarly long-lived and that these long-lived PCs are the major source of serum antibodies; however, only recently have antigen-specific BM PCs been ascertained for their contribution to the pool of serum antibodies in humans^5,12^. Despite these notable advances, the availability of corresponding molecular data (namely, sequence data of BM PC Ig transcripts) and of information regarding PC dynamics *in vivo* are scarce. Alternative models have proposed a role for persistent antigen or else the compartment of memory B cells in the maintenance or renewal of lifelong serological memory, which, in contrast to the notion of intrinsic PC longevity, would allow for continual clonal replacement of antigen-specific PCs^13–15^.

Three studies have generated BM PC data using next-generation sequencing techniques, but none of which have examined the temporal changes that occur in the antibody repertoire over time^5,16,17^. Here, building upon our prior experiences with the comprehensive analysis of human cellular and serological antibody repertoires^18–22^, we present the first longitudinal study of serially acquired human BM PCs assayed by next-generation deep sequencing. To directly measure the temporal dynamics of BM PCs— and to indirectly gain insight into long-lived serological memory—we sequenced recombined VHDJH regions (cDNA), which encode the variable domain (protein) of antibody IGH heavy chains. Most of the VHDJH genetic diversity resides within the CDR-H3 hypervariable interval (encoded by a D element, random non-templated nucleotides, and small portions of the VH and JH elements) and CDR-H3 is a primary determinant of antibody specificity^23,24^ and has long been considered a unique “fingerprint” which aids identification of a progenitor B cell and its clonal progeny (B-cell clonotype)^25^. We sequenced BM PCs from the same individual over seven timepoints encompassing a total of 6.5 years and from a second individual with two timepoints over 2.3 years. The temporal resolution and duration of sampling provides a method to interrogate the *in vivo* temporal dynamics of BM PCs in a previously uncharacterized way. We provide detailed temporal information on the individual genes (IGH V, D, and J), gene combinations (V-D, V-J, D-J, V-D-J), and temporally persistent CDR-H3 clonotypes. The second individual provides support that our observations are not unique. Moreover, persisting CDR-H3 clonotypes were class-switched and somatically mutated (in the IGHV gene segment) implying derivation from activated B cell progenitors that must have been selected by antigen. Crucially, persisting CDR-H3 clonotypes were found exclusively in the PC compartment, but were absent among comparable memory B cells (also class-switched and somatically mutated B-cell compartment) isolated from the same BM biopsy. Overall, our results (i) underscore the temporal stability of the IGH V region repertoire according to multiple metrics (temporally stable IGH molecular phenotypes), and (ii) provide unequivocal sequence-based evidence for the persistence of PC cellular clonotypes spanning 6.5 years.

## RESULTS

### Serial Bone Marrow Biopsies Followed by Next-Generation Sequencing Demonstrate Temporal Dynamics of the plasma cell (PC) Repertoire

To investigate the temporal dynamics of the IGH antibody gene repertoire of bone marrow plasma cells (BM PCs), we sampled, sorted, and performed high-throughput sequencing (Fig. 1a). Serial bone marrow biopsies were obtained from two adolescents (Supplementary Table 1) as part of routine evaluations for non-immuno-hematological disease. BM PCs were isolated using fluorescence-activated cell sorting (FACS). BM PCs were obtained by sorting for CD38++ CD138+ cells within the mononuclear light-scatter gate (Fig. 1b). Additionally, the cells were uniformly positive for the TNF-receptor superfamily member CD27 (Fig. 1b, inset). Importantly, we avoided gating of the pan-B cell marker CD19 since previous characterizations of human BM PCs show heterogeneous expression of CD19^26,27^. Therefore, our method captured all recently described BM PC subpopulations^5,12^ with an overall CD19^+/−^ CD27^+^ CD38^++^ CD138^+^ phenotype. Subsequently, transcripts were amplified from BM PCs expressing IgM, IgG, and IgA using RT-PCR followed by high-throughput sequencing.

**Figure 1.**
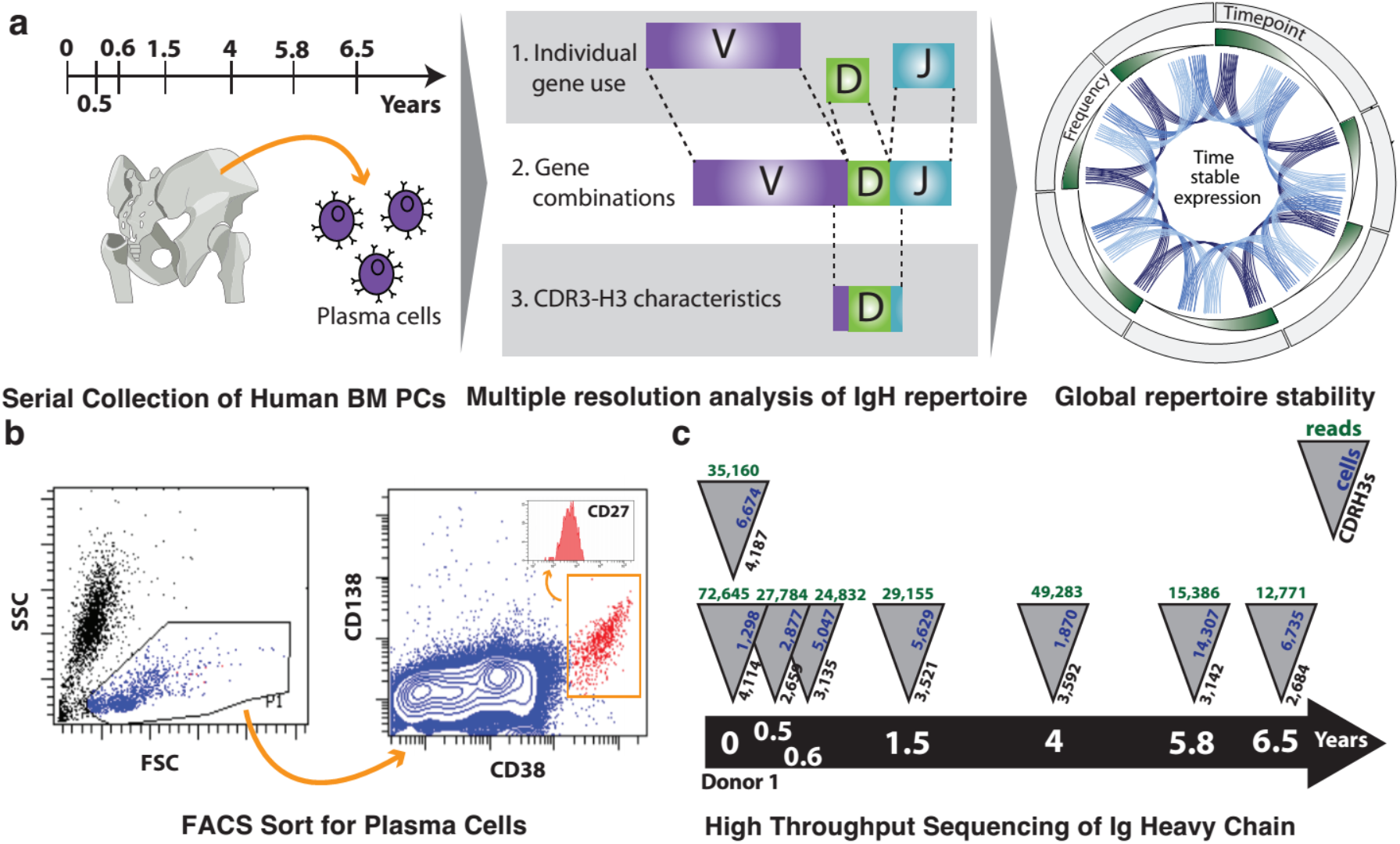
Serial sampling, isolation by FACS, and NGS of bone marrow plasma cells (BM PCs) reveal temporal dynamics of antibody repertoire. (a) Overview of antibody repertoire characterization method. Serial sampling of human bone marrow plasma cells (BM PCs) over 6.5 years (left). Analysis of individual genes, gene combinations, and CDR-H3s (center) show temporally stable expression of persistent entities (right). (b) Representative fluorescence-activated cell sorting (FACS) gates of BM PCs (CD138+, CD38++) isolated from bone marrow mononuclear cells (BMMCs). (c) Sample collection timeline and summary of cell counts, sequencing reads, and unique CDR-H3s.

In total, 51,200 BM PC were sorted, which generated 503,415 reads after quality-threshold filtering (see Methods and Supplementary Table 1). These data were distributed across seven timepoints spanning 6.5 years (Fig. 1c). A biological replicate, a second frozen ampule derived from the same bone marrow aspiration, was also collected from each donor and analyzed. Multiple sampling from the same donor allows us to accurately identify the active heavy chain genes that compose this donor’s antibody repertoire. Specifically, we identify 38 IGHV genes, 21 IGHD genes, and 6 IGHJ genes (4,788 combinations).

### IGH V, D, and J Frequency in the PC Compartment is Highly Stable Over 6.5 Years

To determine the stability of individual gene usage, we assessed the frequency of each IGH V, D, and J gene across time (Fig. 2). Surprisingly, we saw stable behavior of these genes, with the most frequently used genes (e.g. IGHV4–34) showing consistently high expression while less frequently used genes (e.g. IGHV3–72) showed consistently low expression. This observation was quantified using the Mann-Kendall Test, which evaluates trends in time series data. We found that 89% of IGHV genes, 95% of IGHD genes and 100% IGHJ genes showed no statistically significant trends (Mann-Kendall test, p>0.05), indicating that the IGHV (Fig. 2a), IGHD (Fig. 2b), and IGHJ (Fig. 2c) genes were time stable.

**Figure 2.**
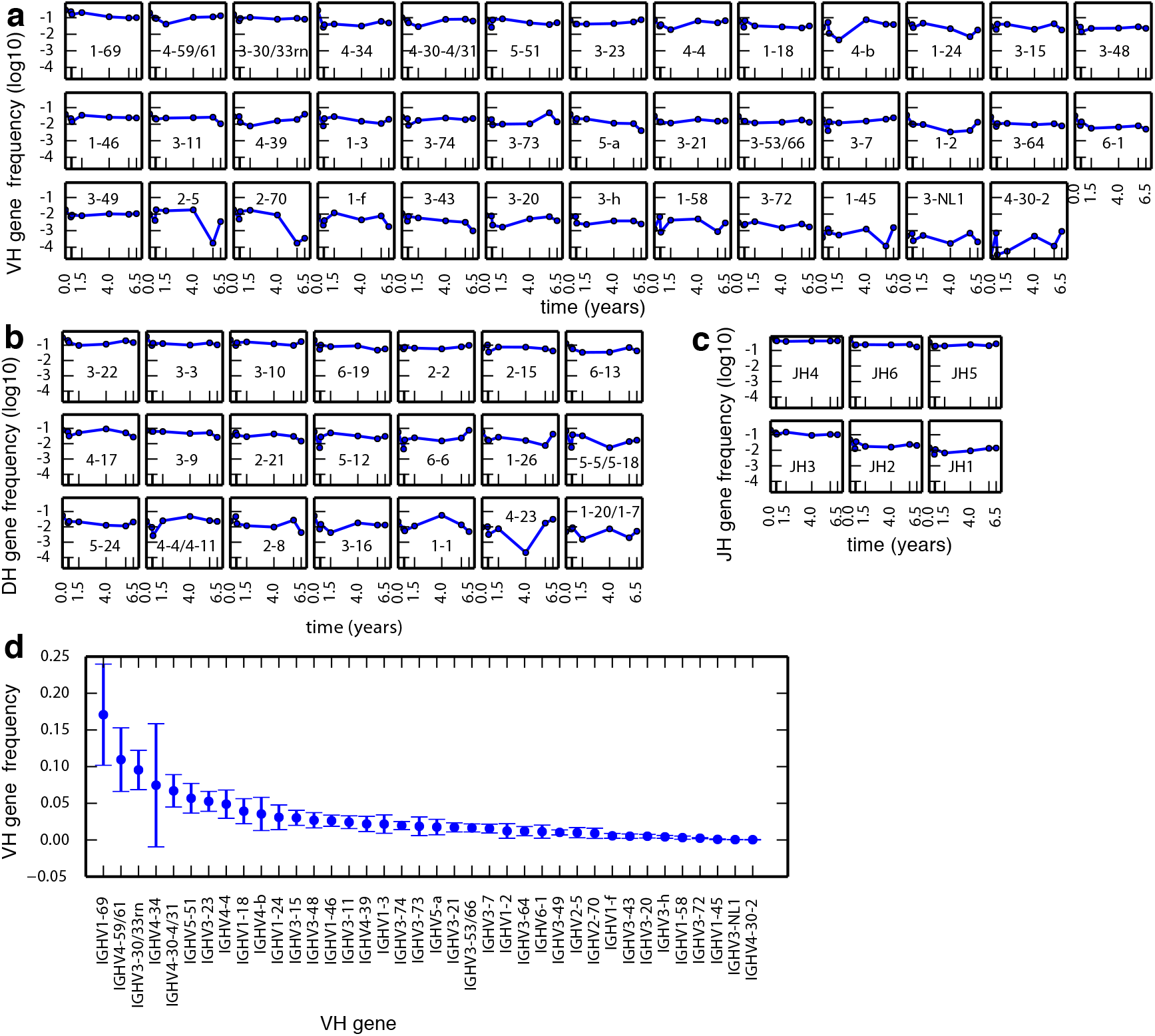
IGH gene segment frequencies among BM PCs are temporally stable. (ac) IGHV (a), IGHD (b), and IGHJ (c) gene usage frequency over time. Plots are sorted by decreasing mean frequency. Only gene calls that appear in all timepoints are shown. (d) Mean frequency of IGHV gene use. Error bars are standard deviation.

Next, we analyzed population behavior of gene usage. Averaging across all timepoints, we observed a highly skewed distribution of individual gene frequencies, consistent with previous single timepoint observations. Only 6 IGHV genes (16%) accounted for greater than 50% of total IGHV gene usage by frequency (Fig. 2d). IGHD genes that have previously been shown to have biased usage IGHD2–2, IGHD3–3, and IGHD3–22^28^ together accounted for 33% of total IGHD usage (Fig. 2b). In addition, known biases in IGHJ usage^29^ are recapitulated as IGHJ4, IGHJ6, and IGHJ5 account for 86% of total IGHJ usage. Furthermore, our analysis demonstrated that IGH V, D, and J gene usage were not significantly different from a log-normal distribution (Anderson-Darling, H=0, p>0.05).

### IGH V-D, D-J, V-J, and V-D-J Combinations in the PC Compartment Are Stable Over Time

Given the temporal stability of individual genes, we hypothesized that differential intrinsic longevity might be found in gene combinations. Surprisingly, our analysis suggests that gene combinations, like their individual component genes, are time stable as well. We found that 92% V-J (Fig. 4), 97% V-D (Supplementary Fig. 1a), 95% D-J (Supplementary Fig. 1b), and 97% V-D-J (Supplementary Fig. 1c) do not show significant trends (Mann-Kendall, H=0, p>0.05).

**Figure 3.**
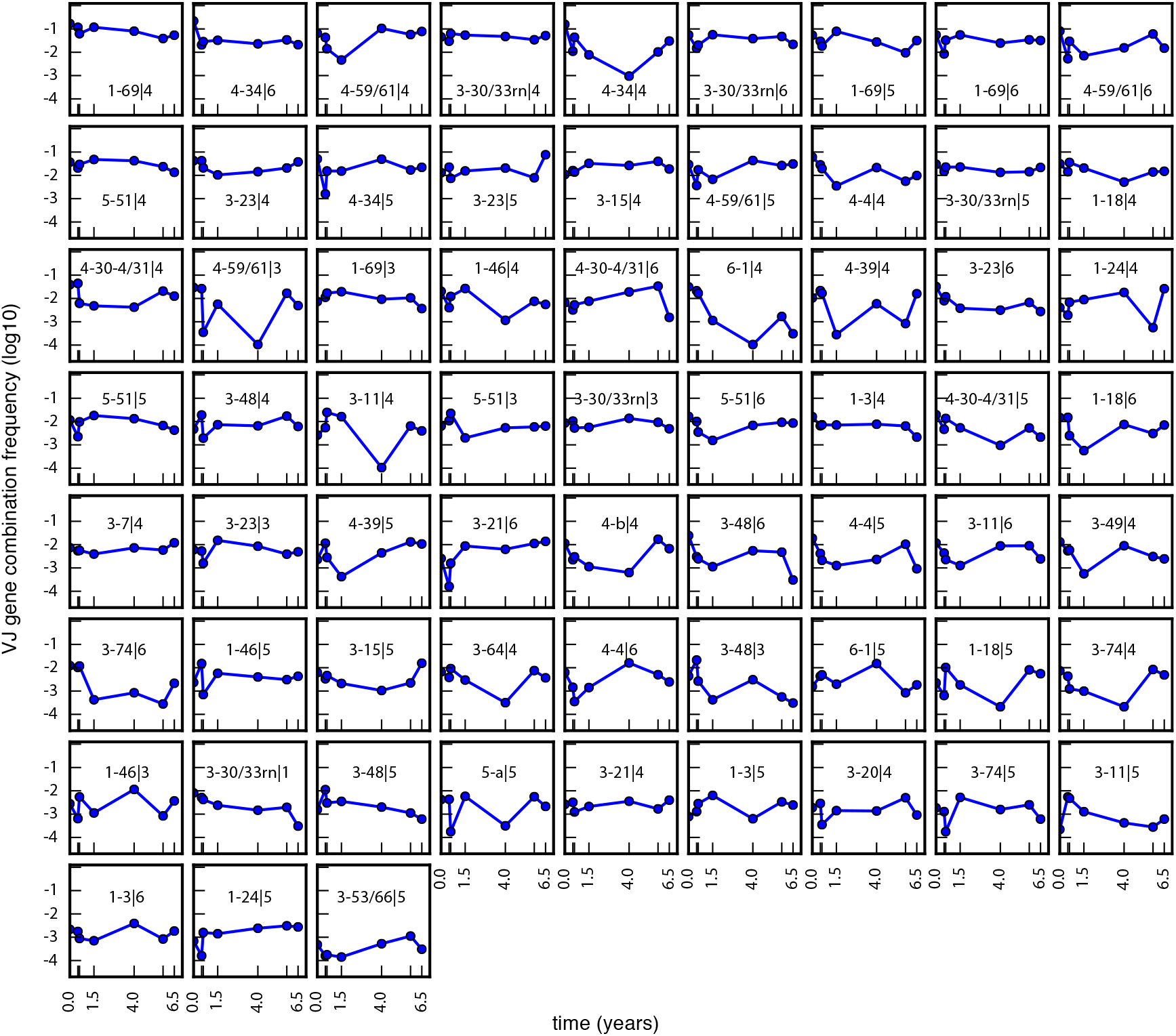
Frequencies of gene combinations among BM PCs are temporally stable. IGH V-J usage frequencies for Donor 1 are shown. Plots are sorted by decreasing mean frequency. Only gene calls that appear in all timepoints are shown. See Supplementary Figure 1(a-c) for usage frequencies of IGH V-D, D-J, and V-D-J.

**Figure 4.**
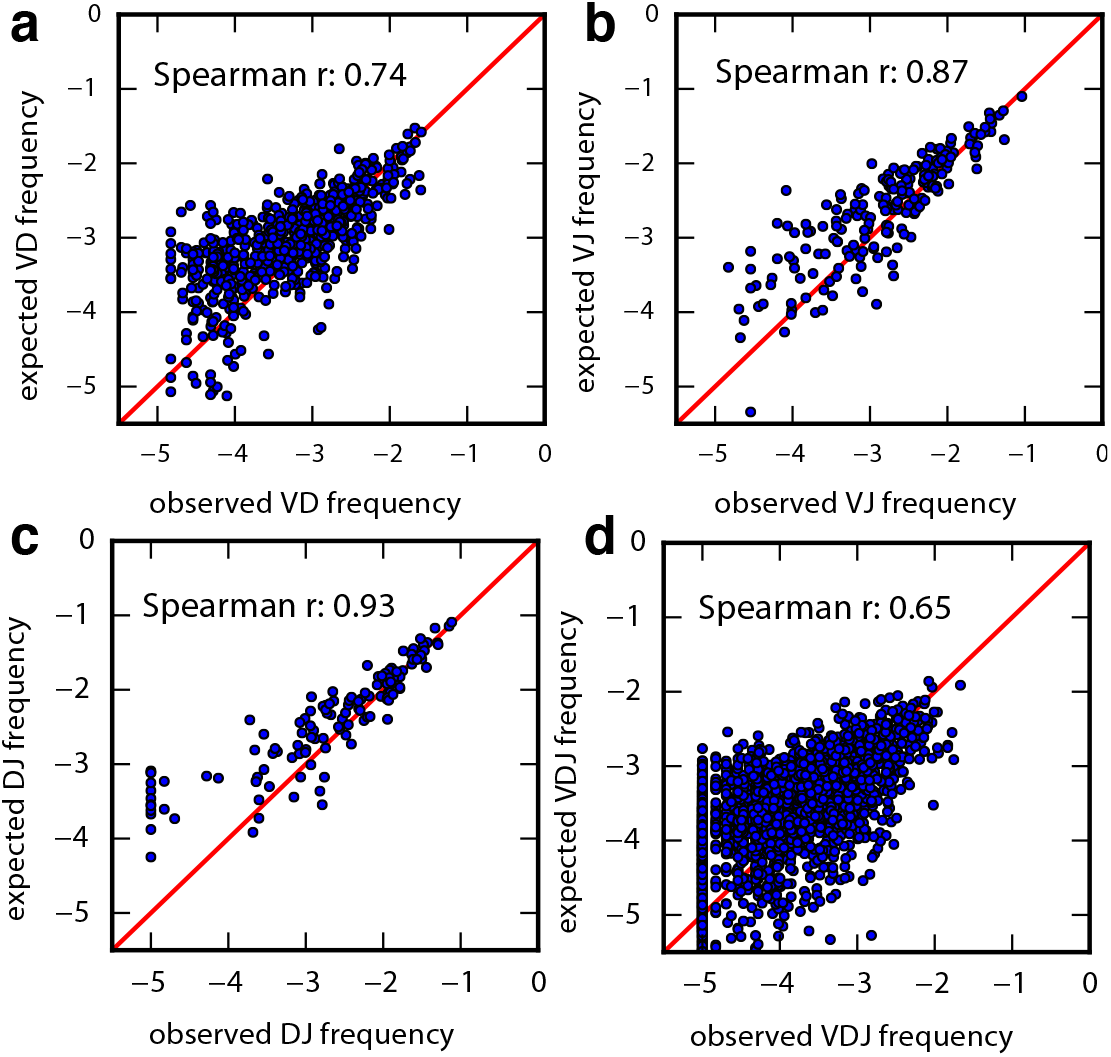
Gene combinations do not preferentially associate and are instead randomly assorted. (a-d) Spearman’s rank correlation of expected versus observed IGH V-D (a), V-J (b), D-J (c), and V-D-J (d) gene combination frequencies. Expected (by random association) frequencies are calculated as products of the frequencies of the individual component genes. Diagonal lines in red indicate no difference between the expected and observed frequencies.

To better understand the nature of gene combinations, we analyzed preferential gene pairing biases by comparing the expected versus observed frequency of pairwise gene combinations. The observed frequency of each gene combination was found to be correlated to its expected frequency (Spearman r): V-D (0.74), V-J (0.87), D-J (0.93), and V-D-J (0.65) (Fig. 5a–d). This high level of correlation and lack of significant outliers suggests that there is minimal gene pairing linkage and that the gene pairing is a random process.

**Figure 5.**
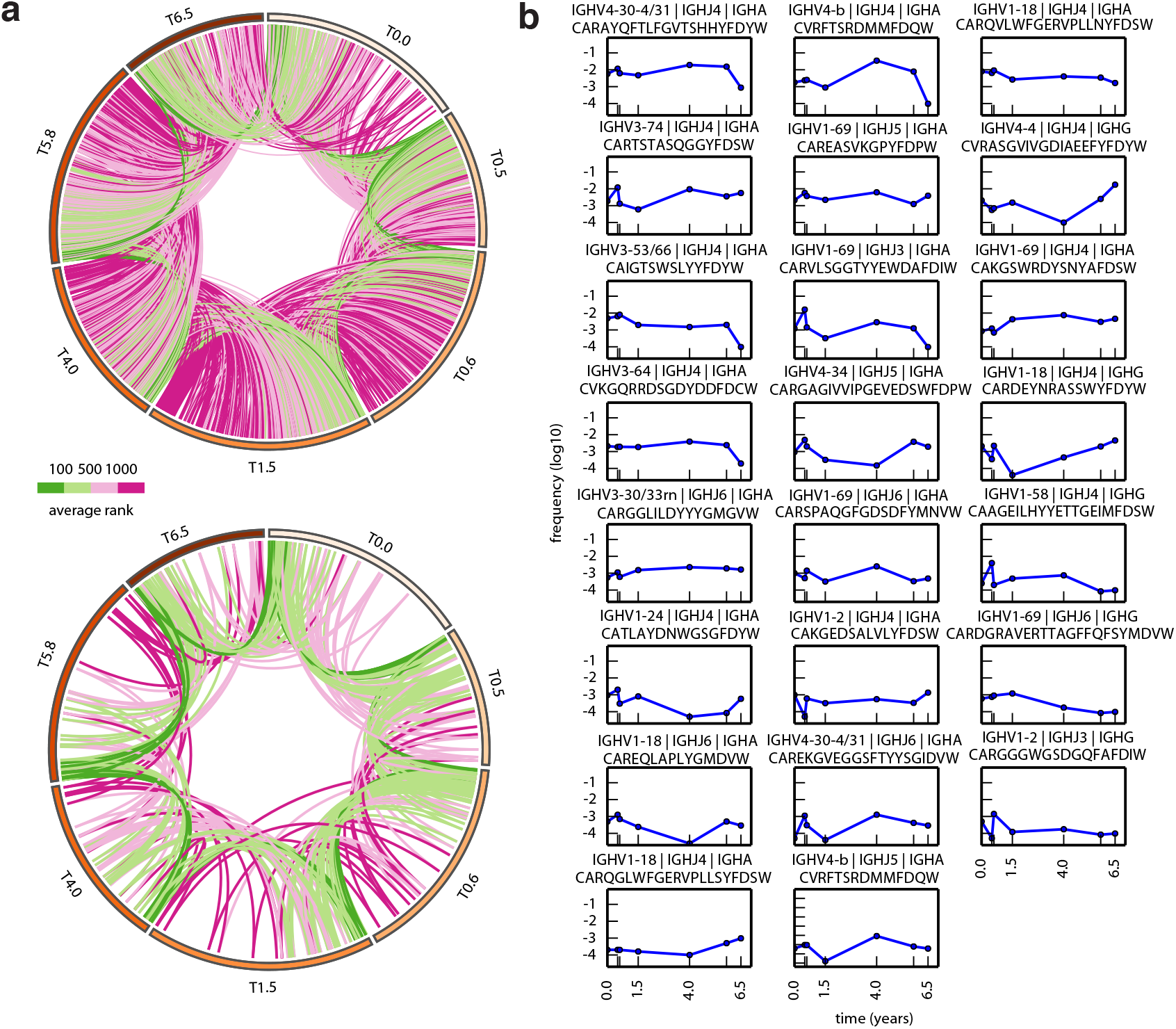
Frequencies of persistent antibody clonotypes among BM PCs are temporally stable. (a) Circos plot of shared CDR-H3 antibody clonotypes between adjacent timepoints across 6.5 years (top). Circos plot of the persistent clonotypes across all timepoints (bottom). Each band in the outermost perimeter represents the clonotypes found in a given timepoint, sorted by decreasing expression. The inner curved lines indicate the same clonotype shared by two timepoints. Green indicates high expression; purple, low expression; with lighter colors indicating intermediate expression. (b) Gene usage frequency over time of the 26 persistent clonotypes (see Methods) found in all timepoints. Plots are sorted by decreasing mean frequency. Gene names (for IGHV and IGHJ), representative amino acid sequences, and isotype are above each plot. (The 153 persistent CDR-H3 antibody clonotypes for Donor 2 are shown in Supplementary Fig. 10)

### Persistent CDR-H3 Clonotypes Are Stable Over Time and Are Unique to PCs Among B-cell Subsets in Bone Marrow

To understand how each of these individual genes and gene combinations together might indicate the existence of long lived PCs, we analyzed the behavior of the CDR-H3, the highest resolution possible for a single identifier of an antibody producing cell. To eliminate errors and ambiguities, we clustered CDR-H3s into clonotypes based on previously established criteria (see Methods). On average, we found that 16% of clonotypes are shared between adjacent timepoints (Fig. 6a, top). Comparison of the BM PC compartment with memory B cells (mBCs) co-isolated from the same biopsy specimens provided a baseline by which to gauge stability across the larger framework of the B-cell compartment: In mBCs, although gene stability (use and frequency) was statistically similar to the PC compartment (Supplementary Fig. 9), not a single persistent CDR-H3 clonotype could be found among 58,953mBCs from the same biopsies across four years in this same donor (data not shown).

**Figure 6.**
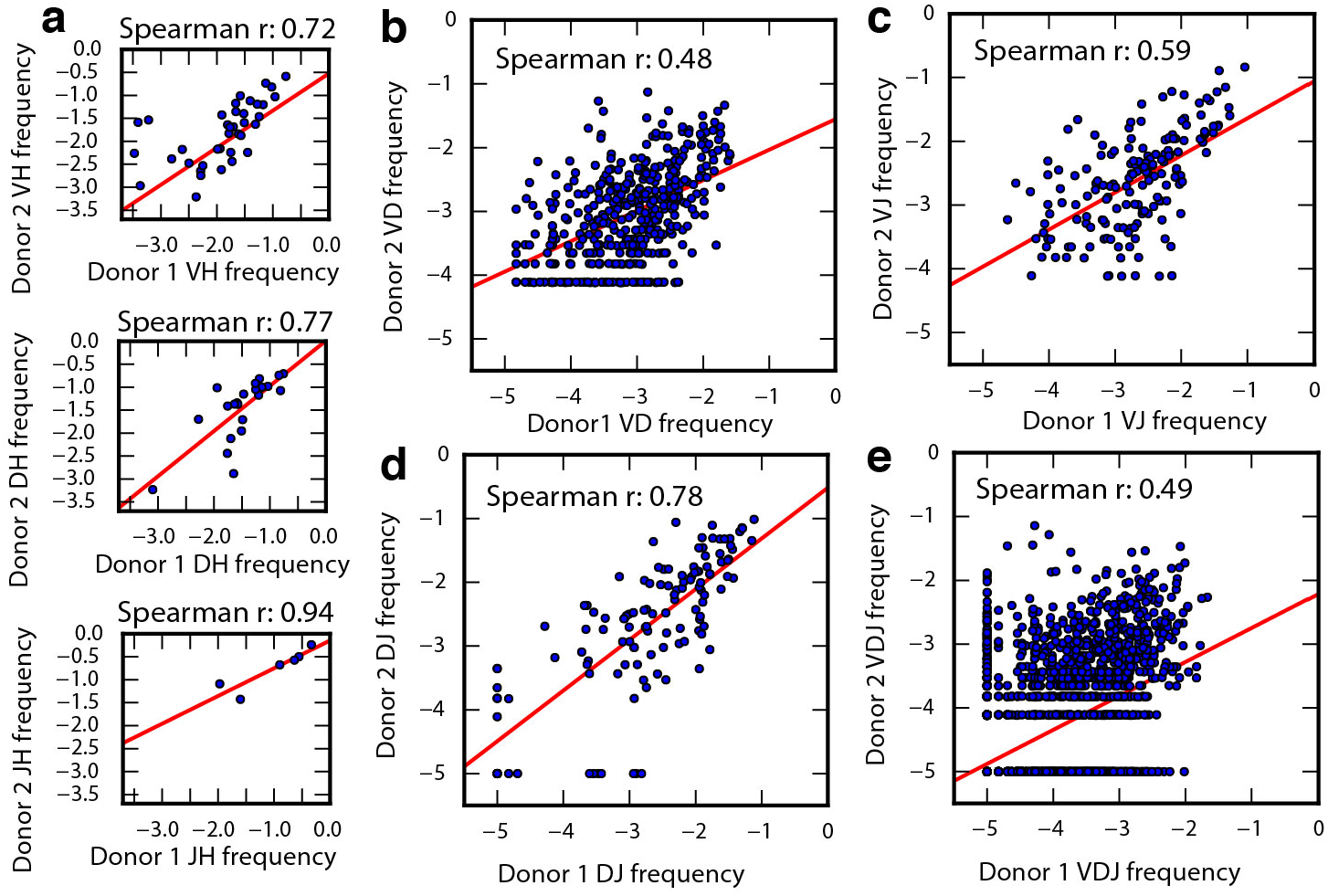
Gene and gene combination use frequencies correlate between Donor 1 and Donor 2. (a) Spearman’s rank correlation of individual gene frequencies between the two donors: IGHV (top), IGHD (center), and IGHJ (bottom). (b-e) Spearman’s rank correlation of combination gene frequencies between the two donors: V-D (b), V-J (c), DJ (d), and V-D-J (e). (a-e) Red lines indicate least squares regression.

Interestingly, among BM PCs, 26 clonotypes persisted across all timepoints spanning 6.5 years (Fig. 6b). We found that 100% of these persistent clonotypes were time stable (Fig. 6a, bottom, Mann-Kendall test, h=0, p>0.05) and 78% (18/23) were of the IgA isotype. In addition, characteristics of the complete CDR-H3 population, specifically CDR-H3 lengths (Supplementary Fig. 2) and hydropathy index (Supplementary Fig. 3), are unchanged over time. The overall total distribution of CDR-H3 lengths are consistent with previously reported single timepoint values. Also, higher expressing CDR-H3s tended to be neither hydrophobic nor hydrophilic (Supplementary Fig. 3) and no significant trends between hydrophobicity and expression level were found.

### A Second Donor Corroborates The Observations From The First Donor

To verify our longitudinal observations of stability and random gene choices from Donor 1, we analyzed a second donor across two years (Fig. 7). We identified 38 IGHV genes, 22 IGHD genes, and 6 IGHJ genes (5,016 combinations, 6,763 cells, 48,525 reads) (Supplementary Table 1 and Supplementary Fig. 4). Donor 1 and Donor 2 show highly correlated IGHV gene usage (r=0.82). Thus, the trends observed in Donor 1 were also observed in Donor 2. Specifically, individual IGHV, IGHD, and IGHJ gene usages were time stable (Supplementary Fig. 4), as were the gene combinations (Supplementary Fig. 5). Consistent with Donor 1, Donor 2 showed no preferential pairing in gene combinations (Supplementary Fig. 6). These results are highly consistent with the trends observed in Donor 1, and together, they indicate that BM PC antibody gene and gene combination usage show surprisingly minimal variation between individuals and across time.

Like Donor 1, no persistent CDR-H3 clonotypes could be found among 24,287 mBCs sorted from the same biopsies across 2.3 years in Donor 2 (data not shown). In contrast, persistent CDR-H3 clonotypes (153) were readily detected in the BM PC compartment (Supplementary Fig. 10). Importantly, these 153 clonotypes were exclusive to the PC compartment (i.e., absent among mBCs). Lastly, as a measure of the quality and integrity of the B-cell sequence datasets derived from the two donors, we observed no inter-donor sequences shared between their mBC compartments, as expected, and only one of the total 179 persistent PC clonotypes was common between the two donors.

## DISCUSSION

Next-generation sequencing has enabled unprecedented ability to explore the details of the human B-cell repertoire^30,31^. Whereas previous studies have been able to describe some aspects of the B-cell repertoire at a single point in time, our results harness the power of next-generation sequencing and longitudinal biopsies of bone marrow (BM) to elucidate the temporal dynamics of BM plasma cells (PCs) over 6.5 years. Importantly, our data provide molecular resolution of antibody identity in the form of CDR-H3 clonotypes, which is not possible with classic techniques like enzyme-linked immunosorbent assay (ELISA).

In this study, we show that the human PC compartment is naturally polarized in both IGH gene choice and gene combination and that the polarization is maintained over time. We also found that the bias is not primarily a result of gene linkage, suggesting there are additional genomic or extrinsic factors that contribute to polarization. As regards genomic factors, next-generation sequencing of identical twin pairs^32,33^ has revealed clear trends for genetic, or heritable, determinants of IGH gene segment use; nonetheless, the CDR-H3 region maintains hypervariability and “fingerprints” interindividual variation that distinguishes twin pairs. Alternatively, the long arms race between the human immune system and the antigens it has confronted throughout evolutionary history may have established a preferential gene choice long ago, and thus the existence of common antibody-mediated solutions to protective immunity; hence, higher expressing genes are likely broad-spectrum antibodies that have been useful in fighting particular classes of disease and continue to do so today. For example, IGHV1–69 is repeatedly implicated in next-generation studies of anti-viral antibody repertoires (e.g., influenza and HIV-1), and the “inherently autoreactive” IGHV4–34 element is associated with a range of autoimmune disorders (e.g., cold agglutinin disease and systemic lupus erythematosus). Indeed, convergent, or “public”, responses using these IGHV gene segments coupled within homologous CDR-H3 clonotypes continue to be newly discovered^34–36^.

Immunological memory is a well-established concept, and memory B cells (mBCs) and BM PCs are thought to be key contributors, in part, through their putative cellular longevity and hypothesized capacity for self-renewal. An outstanding question remains how intrinsic longevity might be established and maintained. It has been proposed that mBCs generate PCs for the lifetime of the human host, and it is further hypothesized that mBCs are endowed with a stem cell–like capacity for self-renewal and as such could be the basis for the continual production of PCs.^37^ Evidence in support of this hypothesis included the demonstration that polyclonal activation of mBCs results in their differentiation into PCs *in vitro*.^14^ As class-switched mBCs coexist with PCs in human bone marrow^38^, we sequenced both compartments to test the hypothesis that mBCs present in the bone marrow might be a renewable source of PCs and, indirectly, the source of long-term Ab production in humans. Because we observed steady usage of IGH gene and gene combinations in both donors throughout our experiment, our observations suggest that there are large resident pools of PCs of the same identity, from which we can sample continuously with no loss of relative expression levels. Moreover, and most importantly, we observed years-long temporal persistence of 179 unique, highly diverse CDR-H3 clonotypes exclusively within the PC compartment, whereas CDR-H3 temporal persistence was devoid in the mBC compartment. This provides a crucial point of comparison between these two B-cell subsets pivotal to immunological memory. The data imply that the molecular sequence stability in the PC compartment is due to persistence of the cellular clonotype, and is not simply a reflection of the naïve B-cell repertoire nor mBC repertoire in general (i.e., heritable influences of IGH gene use, nor replenishment from the mBC compartment).

Whereas ample data has established the persistence of antigen-specific serum Ig titers, which have half-lives of decades or longer^4^, there is to date no insight as to whether the molecular composition of these antibody titers is a homogenous pool of immunoglobulin maintained by a handful of long-lived PC cellular clonotypes or is rather a continual flux and turnover of transitory PC clonotypes. Although our results are unable to verify the lifespan of any one particular plasma cell, we can conclude that clonal members of the CDR-H3 clonotype, which defines identity and binding specificity at the molecular level, does persist for at least 6.5 years. Our results suggest that clonotype persistence contributes to the mechanism underlying long term immunological memory.

In conclusion, we have used high-throughput, next-generation sequencing to definitively identify long-term persistent BM PC clonotypes, which has implications in clinical intervention studies, vaccines, and immunotherapy. Future next-generation sequencing studies can provide an even more detailed picture of the B cell immune repertoire as these studies could include advances in VH:VL native-pair sequencing (paired BCR-Seq^22,39^), the analysis of correlations between BM PC repertoires and serum immunoglobulin species (Ig-Seq^20,21,31^), and an examination of the connectivity of B cells at various developmental stages (e.g., clonal relationships between circulating memory B cells and sessile BM PCs). Our study provides a foundation upon which these further studies can be built.

## MATERIALS & METHODS

### Bone Marrow Specimens

Serially acquired human bone marrow specimens were collected from two donors by aspiration from the ileac crest, and mononuclear cells were enriched by Ficoll hypaque centrifugation. The two adolescent–teenage donors (10–17 years of age) were originally diagnosed with neuroblastoma but had been asymptomatic and disease-free for many years according to routine bone marrow histology. A complete description of the donors’ past medical history and ages at the time of the multiple time point collections is included in Supplementary Table 1. All procedures were performed per a standard operating procedure at the Memorial Sloan-Kettering Cancer Center and collected according to a longstanding protocol approved by the MSKCC Institutional Review Board. Aspirates were withdrawn from four sites and combined (total of 8–10 mL from 4 sites, 2–2.5 mL per site) drawn from the following: anterior right iliac crest, anterior left iliac crest, posterior right iliac crest, and posterior left iliac crest. The same attending physicians performed these procedures and usually biopsied through the same surgical site each time. De-identified specimens were shipped overnight on dry ice to the University of Texas at Austin.

### Flow Cytometry and Isolation of Plasma Cells (PC)

BM samples were quick-thawed in a 37 °C H_2_O bath and slowly diluted into RPMI-1640 complete medium containing DNaseI (Sigma D 4513; 20 U/mL), pelleted, washed and re-suspended in 2 mL FACS buffer (Dulbecco’s PBS + 0.5% BSA Fraction V). Cell viability was determined using Trypan Blue exclusion and on average was approximately 90% per specimen. After a one-hour recovery at room temperature, BM cells were stained for 30 minutes at room temperature using empirically-determined optimal titrations of monoclonal antibodies: CD38-FITC (HIT2), CD138-PE (B-B4), CD27-APC (M-T271), and CD19-v450 (HIB19). CD19^+/−^CD38^++^CD138^+^ cells in human BM were collected as plasma cells (PC). PCs were observed to be heterogeneous for expression of the CD19 B-lineage marker; therefore, CD19-gating was avoided. CD38^++^CD138^+^ PCs were additionally gated by light scatter properties (FSC v. SSC) to exclude debris, apoptotic cells, and remnant granulocytes. In a subset of bone marrow specimens, memory B cells (mBC) were also collected as CD19^+^CD27^+^CD38^−^CD138^−^. Donor 1 included mBCs at 0, 0.5, 0.6, 1.5 and 4.0 years; Donor 2 included mBCs at 0 and 2.3 years. All cell sorts were performed on a FACSAria flow cytometer. Cells were sorted directly into TRI Reagent for RNA preservation.

### RT-PCR, High Throughput Sequencing of IGH V, D, and J Genes

All methods and reagents were as previously described^18^. Variable genes (recombined VHDJH region, which encodes the V domain) of IGH isotypes IgM, IgG, and IgA were amplified from oligo-dT cDNA and sequenced at high-throughput using the Roche 454 GS FLX technology using titanium long-read chemistry.

### Data Processing, Analysis, and Visualization

All sequence data have been deposited to NCBI SRA under BioProject number PRJNA310043. IGHV, IGHD, IGHJ, and CDR-H3 regions for each read was quality filtered, processed and annotated using the VDJFasta utility described previously^40^. Reference IGHV, IGHD, and IGHJ genes from the international ImMunoGeneTics (IMGT) database were used. Mann-Kendall Tests were performed in Matlab, against the null hypothesis of no trend (alpha=0.05). Spearman r non-parametric correlation analysis was performed in python using the scipy library. CDR-H3 sequences were clustered to form antibody clonotypes, as established previously^25,41^, using full-length VHDJH gene nucleotide sequences. VHDJH genes were grouped into clonotypes based on single-linkage hierarchical clustering, and cluster membership required ≥85% identity across the CDR-H3 amino sequence as measured by Levenshtein edit distance.

Circular visualization plots were created with Circos software v0.67–7^42^ where genes were sorted by expression within each timepoint and connected to adjacent timepoints via colored lines showing their expression levels. All other data visualization was performed using Python and matplotlib.

## Acknowledgements

This work was supported by NSF Graduate Research Fellowship DGE-1110007 (to GCW), HDTRA1-12-C-0105 from DTRA (GG, GCI, EMM), WHO GPEI (GCI), and grants from the NIH, NSF, and Welch Foundation (F-1515) to EMM.

## Authorship contributions

GG and NKC conceived the study. GCW and GCI designed and performed experiments, analyzed data, prepared figures, and wrote the manuscript, under the supervision of EMM. All authors reviewed the manuscript.

## Conflict of Interest

The authors declare no conflict of interest.

